# Direct computations of viscoelastic moduli of biomolecular condensates

**DOI:** 10.1101/2024.06.11.598543

**Authors:** Samuel R. Cohen, Priya R. Banerjee, Rohit V. Pappu

**Affiliations:** Department of Biomedical Engineering and Center for Biomolecular Condensates, James McKelvey School of Engineering, Washington University in St. Louis, St. Louis, MO 63130, USA; Department of Physics, The State University of New York at Buffalo, Buffalo, NY 14260, USA

## Abstract

*In vitro* facsimiles of biomolecular condensates are formed by different types of intrinsically disordered proteins including prion-like low complexity domains (PLCDs). PLCD condensates are viscoelastic materials defined by time-dependent, sequence-specific complex shear moduli. Here, we show that viscoelastic moduli can be computed directly using a generalization of the Rouse model and information regarding intra- and inter-chain contacts that is extracted from equilibrium configurations of lattice-based Metropolis Monte Carlo (MMC) simulations. The key ingredient of the generalized Rouse model is the Zimm matrix that we compute from equilibrium MMC simulations. We compute two flavors of Zimm matrices, one referred to as the single-chain model that accounts only for intra-chain contacts, and the other referred to as a collective model, that accounts for inter-chain interactions. The single-chain model systematically overestimates the storage and loss moduli, whereas the collective model reproduces the measured moduli with greater fidelity. However, in the long time, low-frequency domain, a mixture of the two models proves to be most accurate. In line with the theory of Rouse, we find that a continuous distribution of relaxation times exists in condensates. The single crossover frequency between dominantly elastic versus dominantly viscous behaviors is influenced by the totality of the relaxation modes. Hence, our analysis suggests that viscoelastic fluid-like condensates are best described as generalized Maxwell fluids. Finally, we show that the complex shear moduli can be used to solve an inverse problem to obtain distributions of relaxation times that underlie the dynamics within condensates.

## I. INTRODUCTION

Biomolecular condensates are membraneless bodies that form via spontaneous or regulated phase transitions of multivalent proteins and nucleic acids ^1-5^. Macromolecular phase separation is a major component of the phase transitions that contribute to condensate formation ^3,6,7^. As a result, condensates are defined by the presence of two or more coexisting phases. Each pair of coexisting phases is delineated by a phase boundary giving rise to distinct equilibrium and dynamical interphase properties ^8^. Equilibrium interphase properties refer to differences in concentrations of macromolecules and solutes that are engendered by equalization of chemical potentials across phase boundaries ^8,9^. These concentration gradients also create differences in material properties, which we refer to as dynamical interphase properties. They quantify differences in nanoscale dynamics and micron-scale rheology across coexisting phases ^10-24^. Here, we present details of our recent generalization of the Rouse model ^25^, which enables direct computations of viscoelastic moduli for condensates formed by intrinsically disordered proteins such as prion-like low complexity domains (PLCDs) ^26^.

Systematic measurements have yielded detailed descriptions of phase behaviors of the PLCD of the protein hnRNP-A1, which is also referred to as A1-LCD ^27-29^. The phase behaviors of this system and designed variants thereof feature upper critical solution temperatures ^29^. Accordingly, the A1-LCD plus solvent system is in a two-phase regime for temperatures that lie below system-specific critical temperatures and protein concentrations that are above system- and temperature-specific thresholds known as saturation concentrations (*c*_sat_) ^29^. Across the temperature range, the dense phase concentrations vary minimally for most variants. Away from the critical point, the measured volume fraction of A1-LCD variants and other systems within dense phases was found to be ≈0.4 or lower, implying that roughly 60% of condensates are taken up by solvent ^29-31^. While dense phase concentrations change minimally with temperature or for different variants, the values of *c*_sat_ for temperatures below the critical point can vary by several orders of magnitude. The measured values of *c*_sat_ are influenced by parameters such as the number of aromatic residues known as stickers, the types of stickers (tyrosine versus phenylalanine), the local sequence contexts of arginine residues, the ratio of arginine to lysine residues, the net charge per residue, and the glycine versus serine contents ^29^. The ability to modulate and significantly alter the driving forces for phase separation, manifest across roughly forty different designed variants of A1-LCD, has enabled us to probe how the driving forces for phase separation affect material properties of condensates ^26^.

Using passive and active microrheology aided by optical traps, Alshareedah et al.^26^ measured material properties of a subset of the A1-LCD variants studied by Bremer et al.,^29^. They reported the following findings: A1-LCD and designed variants thereof form dense phases that are viscoelastic materials. The dominance of viscous versus elastic moduli depends on the physical age of condensates, and the aging process is highly sequence-specific. The dominantly viscous materials feature system-specific, frequency-dependent complex moduli. Each system has a characteristic timescale known as the crossover frequency (ω_c_). Above the crossover frequency, elastic moduli (*G*′) are larger than viscous moduli (*G*″), whereas the converse is true below the crossover frequency. This frequency is tunable, and it is lowered by increasing the strengths of stickers. Enhancing the strengths of stickers also leads to an increase in the magnitudes of *G*′ and *G*″ across the entire frequency range that can be probed. Finally, there is a clear inverse correlation between measured intra-condensate viscosities and *c*_sat_ values. This implies that stronger driving forces for phase separation translate into higher intra-condensate viscosities. Overall, the measurements were taken to imply that condensates formed by A1-LCD variants are viscoelastic Maxwell fluids. Importantly, the measured viscoelasticities defy assertions of condensates being purely viscous materials such as Newtonian fluids that form via liquid-liquid phase separation ^14,32,33^. Instead, growing evidence points toward phase separation coupled to percolation as the operative process for condensate formation of many intrinsically disordered proteins and nucleic acids ^3,4,34-43^.

PLCDs and other intrinsically disordered proteins are exemplars of linear flexible polymers ^44^. As a result, their dynamics in dilute solutions and in dense phases should conform to expectations based on the Rouse model ^25^, which describes elastic and flow properties of polymer solutions. These properties are governed by the lengths of polymers, their flexibilities and the spectrum of intra-chain and inter-chain interactions between polymer segments, which will be sequence-specific ^25^. It was shown that the Rouse model can be adapted to analyze equilibrium configurations generated via lattice-based Metropolis Monte Carlo (MMC) simulations of two-phase systems formed by PLCDs ^26^. This analysis, aided by computations of Zimm matrices ^45^, allowed for computations of complex moduli and direct comparisons to system-specific measurements. Here, we expand on the work of Alshareedah et al. ^26^ and lay out the details of how the Rouse model was generalized using graph-theoretic descriptions of polymer configurations in dense phases ^46,47^ to compute and analyze sequence-specific viscoelasticities of biomolecular condensates.

The remainder of the narrative is organized as follows: First, we summarize key aspects of the lattice-based MMC simulations ^48^ that were used to generate accurate descriptions of phase behaviors of A1-LCD and designed variants thereof. Next, we present the main tenets of the Rouse model, highlighting the important contributions of the Zimm matrix. We then introduce two approaches for constructing Zimm matrices by adapting graph-theoretic analyses of MMC simulations ^47^. We then show how the eigenvalues of Zimm matrices can be used to compute complex moduli. We conclude with a discussion that summarizes why condensates are best described as generalized Maxwell fluids, and the broader implications for phase separation coupled to percolation being the relevant physics for describing condensate formation ^3,34^.

## II. MMC SIMULATIONS

The simulations we analyze for the computations of viscoelasticity are those of Farag et al., ^47^. They used LaSSI ^36^, which is a lattice-based simulation engine that relies on MMC sampling and a blend of two versions of the bond fluctuation model ^49,50^. The coarse-grained model of Farag et al. used one lattice bead per amino acid residue. The solvent was pseudo-implicit because a vacant lattice site is occupied by solvent molecules. This approach accounts for solvent entropy explicitly, but the solvent-mediated energetics are implicit. Here, we analyze data generated by Farag et al. for two systems that were designated as WT^+NLS^ and allY. The former is the wild-type sequence of A1-LCD that also features a nuclear localization signal (NLS); in the latter, all aromatic residues of WT^+NLS^ were replaced by tyrosine residues. The simulations use *O*(10^2^) molecules and were performed across a series of different temperatures to map coexistence curves.

Monte Carlo moves on a cubic lattice are accepted or rejected based on the criterion of Metropolis et al.,^48^ whereby the probability of accepting a move is the min[1,exp(-Δ*E*/*k*_*B*_*T*)]. Here, Δ*E* is the change in total system energy of the attempted move and *k*_*B*_*T* is the thermal energy. Total system energies were calculated using a nearest neighbor model whereby any two beads that are within one lattice unit of each other along all three coordinate axes contribute to the total energy of the system. The full set of Monte Carlo moves used to obtain coexistence curves included local moves, multi-local moves, reptation or slithering snake moves, translational moves, and pivot as well as double pivot moves ^47^. Biases introduced into the move sets, which were designed to enhance sampling, were accounted for and obviated to ensure that detailed balance is preserved ^36^. Farag et al.,^47^ introduced multi-local and pivot moves that add to the move sets in the original LaSSI engine ^36^. The multi-local move attempts to move a single bead and its covalently bonded partners by -1, 0, or 1 lattice unit along each coordinate axis. The pivot move chooses a random chain, then chooses a random bead, *x*, along this chain. The move attempts to pivot every bead in the chain beyond *x* in the same direction by 90°.

To calculate coexistence curves, Farag et al., performed LaSSI-based MMC simulations at a series of simulation temperatures with several hundred chain molecules in the simulation setup. All simulations used a 120×120×120 cubic lattice with periodic boundary conditions. The simulations were initialized in a smaller 35×35×35 cubic lattice, and the box size was changed after initialization. This approach allows for significantly faster equilibration of coexisting phases. However, it precludes the analysis of pre-equilibrium processes, such as condensate coalescence. For each variant, 200 chains, each with 137 beads, were placed in the cubic lattice. Thus, the total volume fraction of beads was approximately 0.016.

A total of 1.5×10^10^ steps was used at each simulation temperature. Equilibration was typically realized after about 2×10^9^ steps, as determined by plateauing of the total system energy. All simulation results were analyzed after the halfway point of 7.5×10^9^ steps. Two distinct order parameters are typically computed in LaSSI simulations. Analysis of percolation uses the approach of Harmon et al., ^51^ which quantifies the concentration threshold required for forming the single largest cluster that is system-spanning by comparing it to the percolation threshold computed analytically ^37^. The coexistence curve was computed using the radial density profiles for the distribution of polymer beads in the system, resulting in estimates for concentrations of coexisting dilute- and dense-phases. All simulations were performed using at least three replicates, with each simulation initiated by a distinct random seed. The computed coexistence curves were compared to the measurements of Bremer et al.,^29^ and the agreement was found to be very good ^47^.

We analyzed the dense phases of WT^+NLS^ and allY condensates by extracting snapshots from the LaSSI simulations. The dense phase is taken to be the largest cluster of connected chains. Two molecules are treated as being connected if at least one pair of residues from the molecules in question are nearest neighbors on the cubic lattice, which we define as belonging to any of the 26 sites that surround a given lattice site. This approach is used to quantify the numbers of intra- and inter-chain contacts for molecules in the dense phase.

## III. THE ROUSE MODEL FOR VISCOELASTICITY

The theory of Rouse describes the near-equilibrium dynamics of random coil polymers in the linear response regime ^52^. The equations of motion for the Rouse theory can be expressed in matrix form using a Langevin equation:

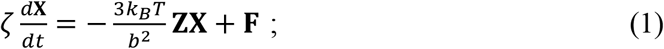

Here, ζis the friction coefficient of the background, *k*_*B*_ is the Boltzmann constant, and *T* is the temperature. **F** is the random force that satisfies the fluctuation-dissipation relation. **X** is the column vector of the positions of Kuhn monomers of length *b*. The pre-factor of 3*k*_*B*_*T*/*b*^2^ comes from treating Kuhn monomers in a bead-spring model (Fig. 1a) with mean end-to-end distances that conform to a Gaussian distribution. A key component of the Rouse model is the Zimm matrix, **Z**. In the Rouse model, this matrix describes the connectivity between Kuhn monomers. Here, we adapt the Zimm matrix to account for the network of physical crosslinks that form between monomers or effective monomers in dense phases formed in equilibrium MMC simulations of two-phase systems. In our approach, we average across the ensemble of conformations for individual molecules or networks of molecules. Accordingly, the Zimm matrix contains all the information regarding the relaxation of a polymer to equilibrium.

**Fig. 1:**
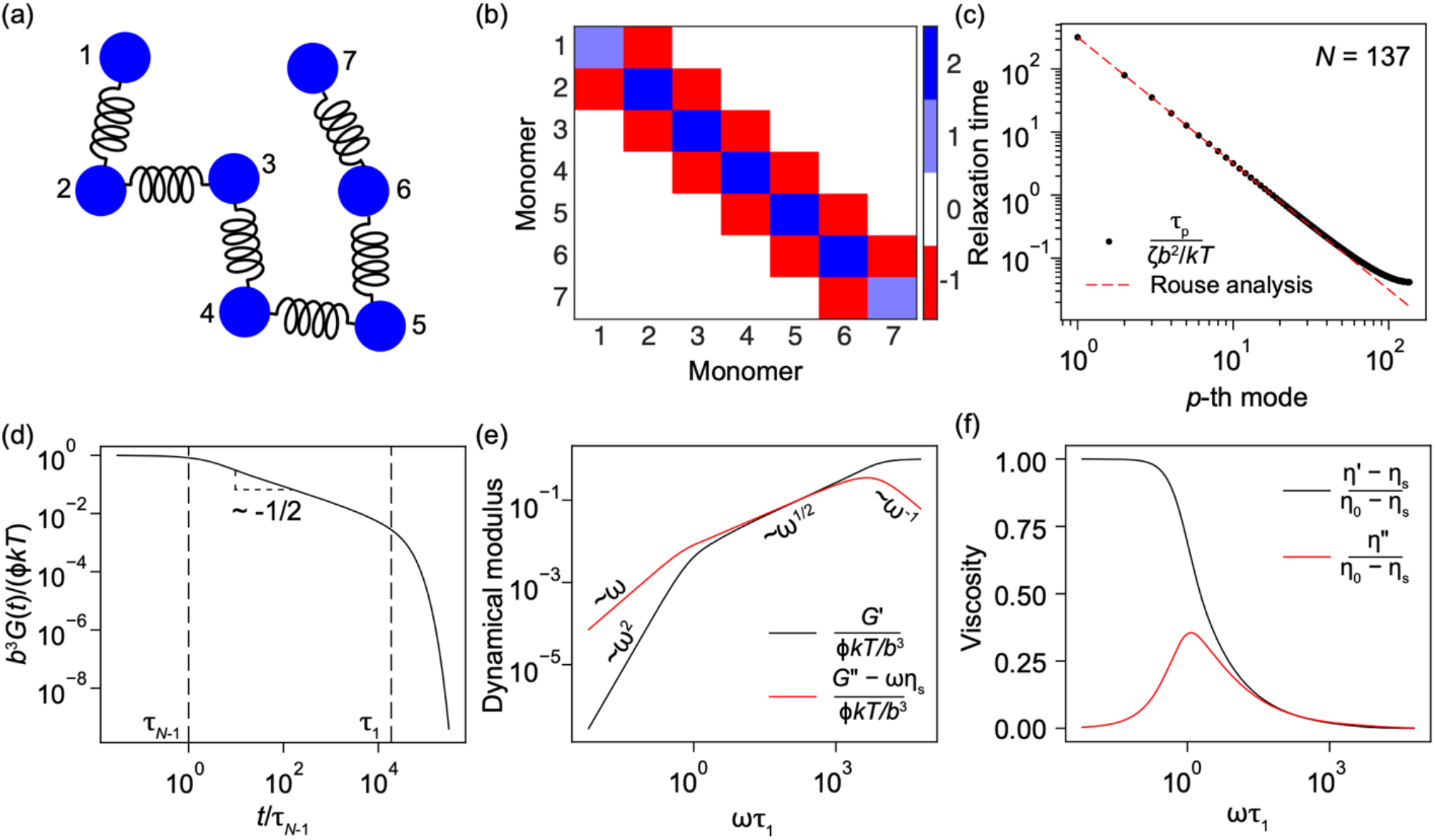
Rouse theory for an ideal chain in solution. (a) Bead-and-spring model for a single chain (*N* = 7). The beads are Kuhn monomers, each of size *b*. (b) The central object of our adaptation of the Rouse theory is the Zimm matrix, which is shown here for an ideal chain of seven beads. (c) The distribution of relaxation times, τ_*p*_, for a 137-mer chain is calculated from the eigenvalues of the Zimm matrix. A fit of the dimensionless relaxation times to [1/(6π^2^)](*N*/*p*)^2^ agrees well with the slowest relaxation times. The mode corresponding to τ_*N*−1_ describes the relaxation time of a single monomer, whereas τ_1_ describes the relaxation time of the entire chain. (d) Plots of the computed relaxation moduli for an ideal 137-mer. (e) The storage modulus (*G*′) and the loss modulus (*G*″) for an ideal chain are plotted against angular frequency. The scaling ∼ω^1/2^ in the frequency range, 1/τ_1_ ≪ ω ≪ 1/τ_*N*−1_, is observed, with *G*″ converging to *G*′. (f) Viscosities are calculated according to *i*ω*η*^∗^ = *G*^/^ + *iG*^//^. The quantities here are normalized to the zero-shear viscosity, *η*_0_, and the polymer-mediated viscosity of the solvent, *η*_*s*_. In (c)-(f), we plot the dimensionless quantities.

We take advantage of the fact that the Zimm matrix is formally equivalent to a graph Laplacian ^46^ wherein polymers or beads within a polymer are nodes that are connected to other polymers or beads by edges. The nodes can be Kuhn monomers or entire chains. The edges are either covalent crosslinks for ideal chains, a combination of covalent and non-covalent crosslinks for a real chain, or non-covalent crosslinks between chains if each node is a single chain. The graph Laplacian is defined as the difference between the degree (**D**) and adjacency (**A**) matrices ^53^ such that **Z** ≡ (**D** – **A)**. In the degree matrix **D**, the off-diagonal elements are zero, and the diagonal elements quantify the number of connections for each node. Elements of the adjacency matrix, **A**, have a value of 1 if two nodes are connected or 0 if they are not connected by either covalent or non-covalent crosslinks. In Fig. 1b we illustrate the Zimm matrix for an ideal linear chain. The graph is undirected and unweighted. Here, the diagonal elements of the Zimm matrix have values of 2 or 1 depending on whether the monomers are internal to the chain or are at the ends of the chain. A value of 2 signifies the fact that each monomer is connected to one bead on each side, whereas the two terminal beads have one connection each. For an ideal chain, all off-diagonal elements other than the band diagonal elements are zero. The band diagonal elements have values of –1.

Eq. (1) can be solved by introducing normal coordinates where the positions are independent of one another ^25,54^. Since the Zimm matrix is a graph Laplacian, we rewrite it as **Z** ≡ **BB**^*T*^, where **B** is the unoriented incidence matrix in graph theory. Accordingly, for an undirected graph consisting of *N* nodes and *m* edges, the elements of the *N* × *m* incidence matrix are set to 1 if there is a node *i* that is incident upon edge *j*; otherwise, the elements are set to zero. We use the incidence matrix to rewrite Eq. (1) as:

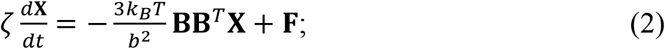

Following the approach of Rouse ^25^, we define a new set of coordinates, **r**, which we can relate to the original coordinates using:

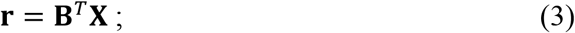

Multiplying both sides of Eq. (2) by **B**^*T*^ gives:

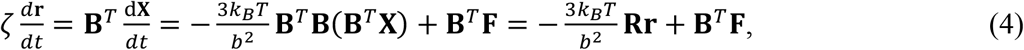

where **R** ≡ **B**^*T*^**B**. If we only consider the first term in Eq. (4), because simulations account for the random force that perturbs the system from equilibrium, the solution to Eq. (1) becomes a solution to an eigenvalue problem:

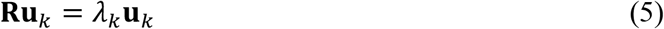

for eigenvectors **u**_*k*_ and eigenvalues λ_*k*_ for *k* = 1, 2, …, *m*, where *m* is the total number of springs in the Rouse model or edges in graph-theoretic instantiations of the Rouse model. The solution to Eq. (1) therefore requires solving for all nonzero eigenvalues of either the *m* × *m* Rouse matrix, **R**, or of the *N* × *N* Zimm matrix, **Z**, which has been shown to be equivalent ^46^. Note that for *N* monomers or nodes, there are *N* − 1 nonzero eigenvalues. This follows from spectral graph theory, where the first eigenvalue of the graph Laplacian is zero for a fully connected graph, such as in the Rouse model of a polymer ^55^.

### A. The Zimm matrix and its importance

We are interested in the eigenvalues of the Zimm matrix:

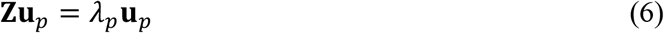

for *p* = 1, 2, …, *N* − 1. Following Zimm ^45^, we diagonalize **Z** using:

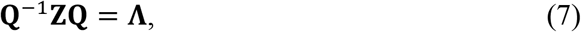

where **Q** is the *N* × *N* matrix with the eigenvectors **u**_*p*_ as its columns, and **Λ** is the diagonal matrix of the eigenvalues λ_*p*_. The matrix **Q** can be used to transform the original coordinates **X** into normal coordinates **q** according to:

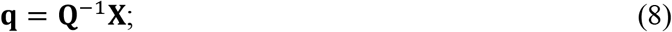

Taking the time derivative of both sides of Eq. (8) leads to:

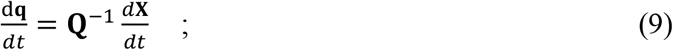

By comparison with Eq. (1) and Eq. (8) we have:

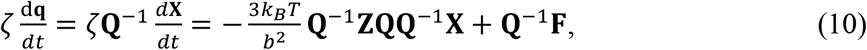

and this simplifies to:

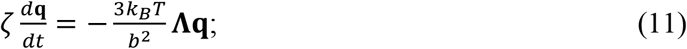

where we have neglected the contribution of the random force since it is not needed to solve the eigenvalue problem ^54^. To arrive at Eq. (11), we use the property that **QQ**^-1^ = **Q**^-1^**Q** = **I**, the identity matrix. The normal coordinates are related to their time derivative by:

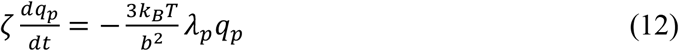

and this yields the expression,

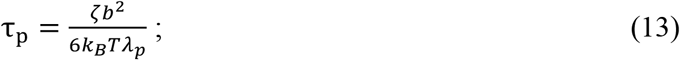

The expression in Eq. (13) defines the full distribution of relaxation times, τ_*p*_ ^54^. Therefore, the distribution of dynamical modes, as captured in the spectrum of relaxation times for an ideal polymer in solution, comes from knowledge of the Zimm matrix. We adapt this feature for real chains and analyze simulations of two-phase systems by constructing Zimm matrices that describe the equilibrium structures within dense phases. Prior to describing this adaptation, we discuss key dynamical properties of an ideal polymer in a solvent that can be extracted by analyzing the distribution of relaxation times.

### B. Results from analysis of relaxation times for an ideal polymer in a solvent

In the Rouse theory, collisions of the Kuhn monomers with the solvent contribute stochastic and dissipative components to the force exerted by the solvent on the chain. Fluctuations due to thermal agitation and the dissipative effects lead to a relaxation to equilibrium configurations, described by Gaussian statistics. The use of normal coordinates, as shown in Eq. (11) resolves the motions of the polymer into a series of modes. Each of the modes is defined by a characteristic relaxation time. For an ideal chain of *N* Kuhn monomers, there are *N*–1 non-zero modes. Since A1-LCD is a 137-mer, we first computed the distribution of relaxation times, τ_*p*_, for an ideal 137-mer chain (Fig. 1c). These relaxation times are inversely proportional to the eigenvalues of the Zimm matrix. The mode corresponding to τ_*N*-1_ describes the relaxation time of a single monomer, whereas τ_1_ describes the relaxation time of the entire chain. A fit of the dimensionless relaxation times to [1/(6π^2^)](*N*/*p*)^2^ agrees well with the slowest relaxation times (Fig. 1c). As noted by Rouse ^25^, this fit should not work for the fastest modes, and we find a clear deviation for the higher *p* modes, which correspond to the dominant contributions made by the elasticity of the ideal chain (Fig. 1c).

Following Rouse ^25^, we use the distribution of relaxation times, τ_*p*_, to compute the complex shear modulus in terms of the storage and loss moduli using:

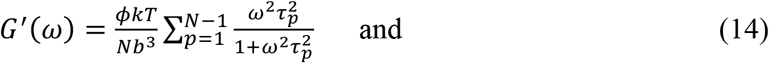

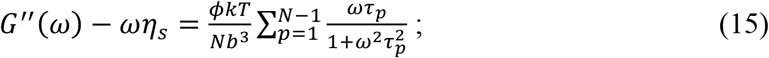

Here, ϕ is the volume fraction of polymers in solution, ω is the angular frequency, and η_s_ is the viscosity of the solvent. We do not need to know the solvent viscosity since we compute the loss modulus dereferenced against (ωη_s_) in Eq. (15), and all quantities on the right-hand side are known. In our computations, we set ϕ, *b*, and ζto be unity, and *T* is in simulation units with *k* set to unity. These choices are made because the quantities are pre-factors that do not materially alter any of the conclusions. Furthermore, when comparing to experiments, we use a single parameter, borrowed from the measurements that put the computed and measured values on the same scale. We use a simulation temperature of 53, which corresponds to 296.8 K for A1-LCD ^28,47^, thus closely matching experimental conditions. Using the same values for the constants, we also compute the relaxation modulus for an ideal chain, which is the time-domain analog of the complex shear modulus:

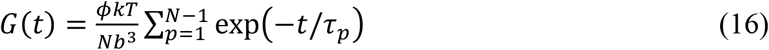

The relaxation modulus scales as 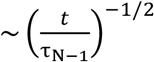 in the regime between τ and τ. Our analysis of the eigenvalues for the ideal chain reveals interesting behaviors for the frequency dependence of the storage and loss moduli. In the low-frequency regime, *G*′ < *G*″, and the storage modulus *G*′ scales as ∼ω^2^ whereas the loss modulus *G*″ scales linearly with ω. In the high-frequency regime, *G*′ > *G*″, and while *G*′ plateaus, *G*″ decreases as ∼*ω*^-1^. Between the regime where *G*′ < *G*″, and *G*′ > *G*″, there is a regime where *G*′ = *G*″. In this regime, 1/τ_1_ ≪ ω ≪ 1/τ_*N*-1_, both *G*′and *G*″ scale as ∼ω^1/2^.

The complex shear moduli show that even an ideal polymer in a dilute solution has finite elastic and viscous moduli. Given the *N*–1 finite relaxation times, Rouse proposed that the viscoelastic properties of ideal polymer solutions conform to a generalized Maxwell model ^25,56^. In this conception, Maxwell elements representing each of the relaxation modes, contribute on the order of *k*_*B*_*T* to the overall storage modulus at high frequencies. Next, we calculate the complex viscosities of the dilute solution of ideal chains using *i*ω*η*^∗^ = *G*^′^ + *iG*^″^. Plots of these normalized quantities reproduce the results of Rouse ^25^.

Next, we asked how the inclusion of non-covalent interactions in the form of contacts between non-nearest neighbors along a linear chain influences the storage and loss moduli of an ideal chain. These additions, which are relatively sparse, are made to ensure that the contacts do not significantly perturb the ideal chain statistics. As shown in Fig. 1c-1f, we consider a 137-mer. For an ideal chain, the off-diagonal density of the Zimm matrix is π=0.0146. This non-zero density is entirely due to bonded interactions. We randomly added non-bonded connections to this system by increasing the density of off-diagonal elements. As random contacts are added, the intermediate region, where *G*′ = *G*″ and the moduli scale as ∼ω^1/2^, shrinks such that the crossover corresponding to the equality of the moduli becomes a crossover point. Adding random non-bonded contacts to an ideal chain yields frequency-dependent profiles for the complex shear moduli that are concordant with a conventional Maxwell model ^57^. The changes to the viscoelastic responses observed by the addition of random, non-bonded interactions motivated computations of Zimm matrices by analyzing contact patterns from the LaSSI-based MMC simulations of phase equilibria for WT^+NLS^ and allY versions of A1-LCD.

## IV. ZIMM MATRICES EXTRACTED FROM MMC SIMULATIONS

Analysis of the LaSSI simulations showed that chains are more expanded within dense phases when compared to coexisting dilute phases ^47^. The concentrations of A1-LCD and related molecules within dense phases are above the overlap regime ^47^. Accordingly, intra-chain and inter-chain interactions are likely to occur with similar probabilities, and the chains have near Gaussian-like character within dense phases. These observations justify our use of the Rouse model, where the main determinant of the complex shear moduli are the eigenvalues of the Zimm matrix. Having established that the Rouse theory can be deployed to account for non-bonded interactions (see Fig. 2), we extracted Zimm matrices from lattice-based MMC simulations of the A1-LCD system (Fig. 3a). The approaches we describe below would need to be modified if globule- or rod-like conformations are dominant in the dense phase ^58^, although recent data suggest that even globular domains undergo some degree of unfolding inside dense phases ^59,60^.

**Fig. 2:**
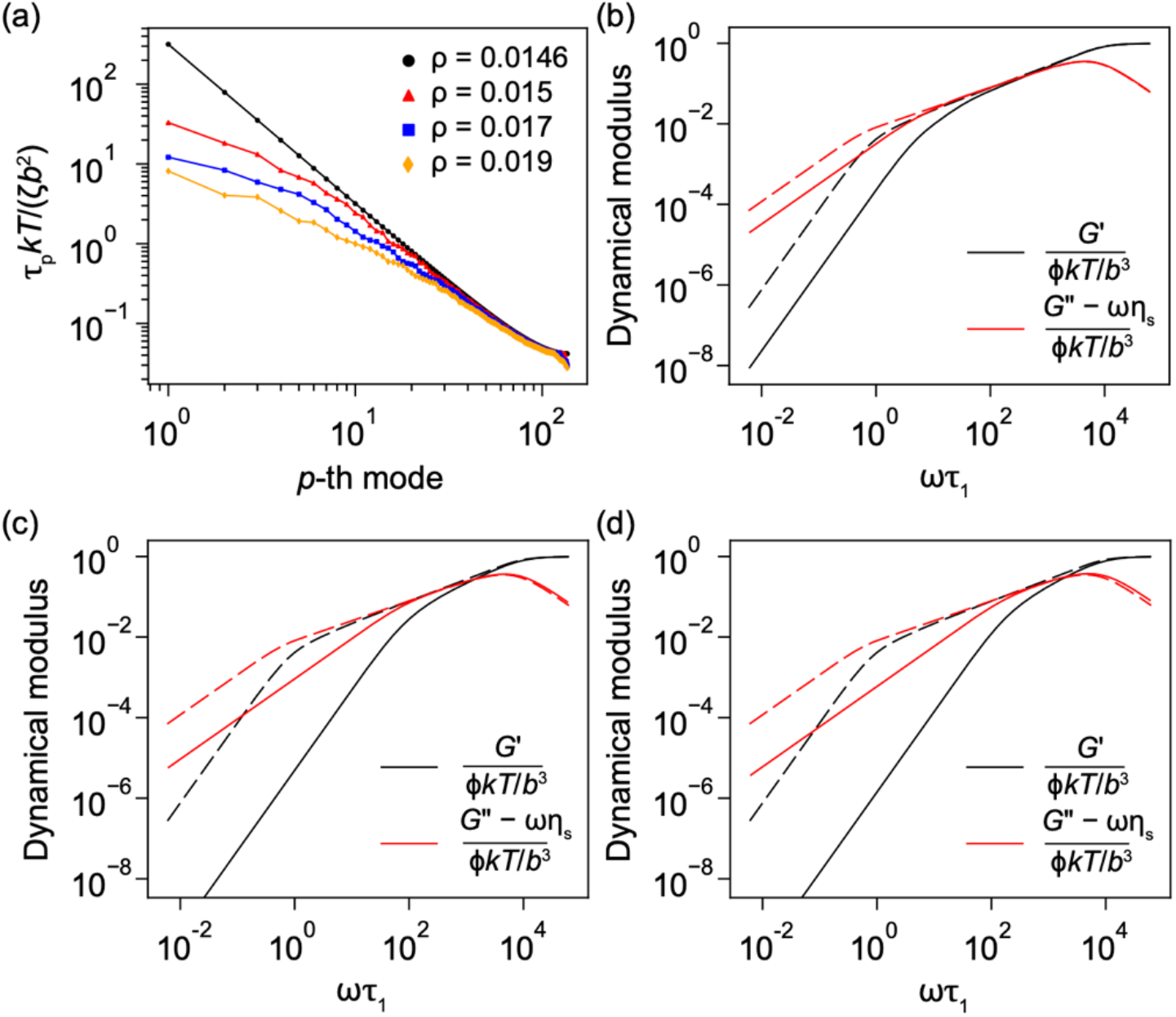
Assessments of how non-bonded contacts impact computed moduli of an ideal chain. (a) Starting with the 137-mer ideal chain, we randomly add contacts by increasing the off-diagonal density of the Zimm matrix starting with the ideal chain at *ρ* = 0.0146. We compute the dynamical moduli (solid lines) and compare the results to the ideal chain (dashed lines) for (b) *ρ* = 0.015, (c) *ρ* = 0.017, and (d) *ρ* = 0.019. The intermediate region in the dynamical moduli disappears as random contacts are added.

**Fig. 3:**
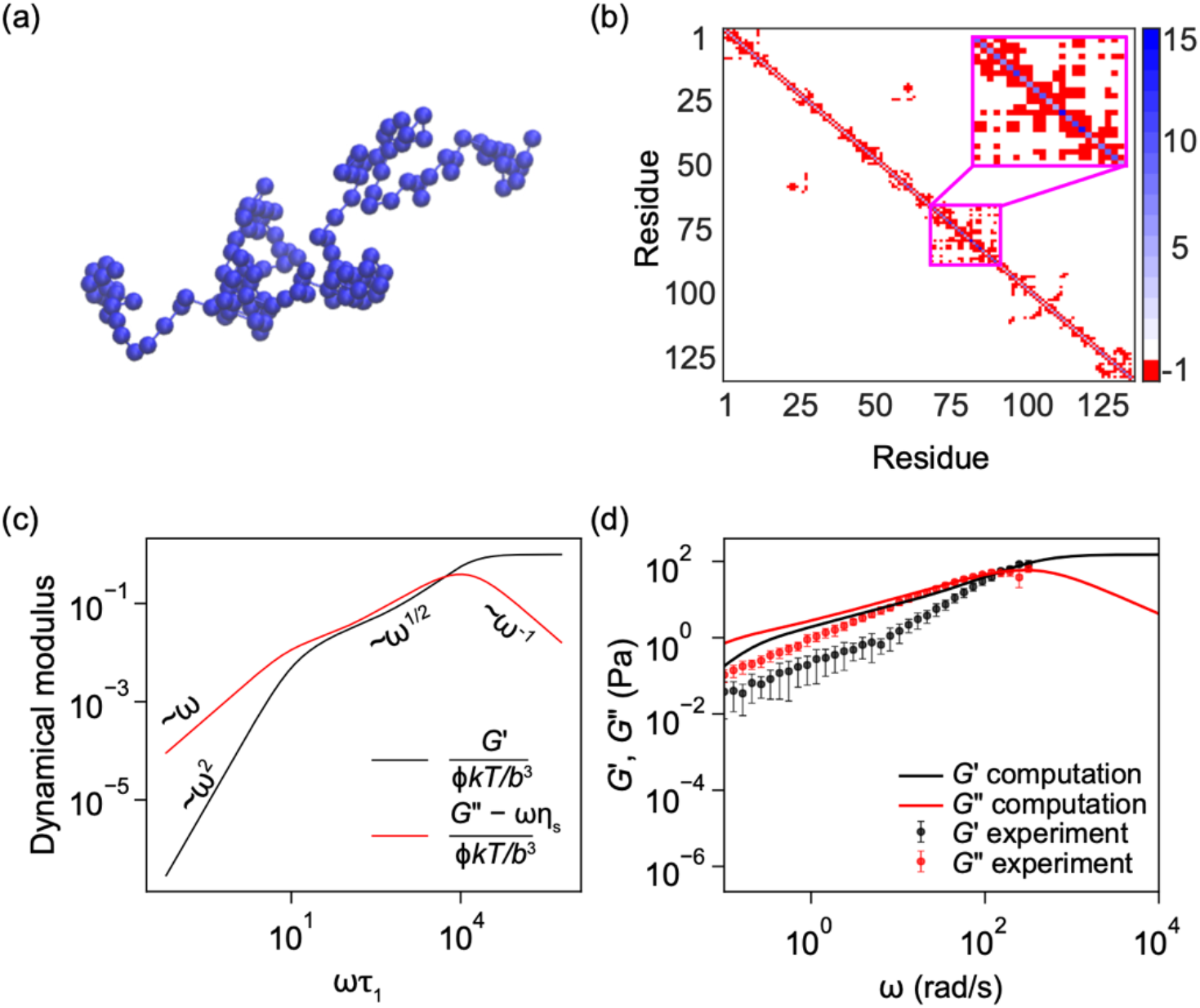
The single-chain model and comparisons of computed and measured moduli. (a) Snapshot of a single chain extracted from MMC simulations of the dense phase of A1-LCD: WT^+NLS^. The residues are considered as Kuhn monomers. (b) The Zimm matrix includes both bonded and non-bonded intra-chain interactions. The inset indicates a region of the Zimm matrix showing the types of intra-chain contacts that form in dense phases. (c) The corresponding dynamical moduli show an intermediate response scaling as ∼ω^1/2^. The Rouse time, τ_1_, is smaller than for the ideal chain due to internal friction imparted by the non-bonded interactions. (d) Comparisons to measurements, following a rescaling to match the measured crossover frequency, show that the single-chain model overestimates the experimental storage and loss moduli. The computed moduli were analyzed over 30 snapshots across 3 replicates by averaging the *p*-th relaxation times over all chains in the dense phase.

### A. The single-chain model

We first tested an approach that we termed the single-chain model (Fig. 3b). Here, the nodes represent residues on a single chain within the condensate, and the edges are defined by bonded and non-bonded intra-chain contacts. The Zimm matrix was calculated as **Z** ≡ **BB**^*T*^, where the unoriented incidence matrix **B** has elements *B*_*ij*_. The elements *B*_*ij*_ are set to 1 if there is a node *i* incident upon edge *j*; otherwise, *B*_*ij*_ equals zero. For *N* nodes and *m* edges, the incidence matrix has size *N×m*. This allows us to consider the effect of edge-weighting the graph. Here, we used the unweighted graphs.

For distinct snapshots in the simulations, chains located within the interiors of dense phases were analyzed for the presence of non-bonded intra-chain contacts. The ensemble-averaged Zimm matrices were then used to compute the eigenvalue spectrum and the complex shear moduli. The frequency dependence of the computed moduli (Fig. 3c) shows clear deviations from the profiles for ideal chains (Fig. 2e). For the single-chain model used to capture the dynamics within a condensate the Rouse time, τ_1---_, which is the slowest relaxation mode, is longer than for an ideal chain in a dilute solution. This is due to internal friction imparted by the non-bonded interactions. However, when compared to what we observe by adding random, non-bonded contacts at higher off-diagonal densities (Fig. 2c and Fig. 2d), the intermediate regime where the moduli scale as ∼ω^1/2^ is largely preserved, although the storage and loss moduli are unequal in this regime. This highlights the sparsity of intra-chain contacts for A1-LCD molecules within the dense phases. The sparsity of intra-chain contacts is readily attributable to the chains forming expanded conformations within dense phases ^47^.

Next, we compared the moduli computed using the single-chain model to the moduli measured using passive microrheology with optical trapping ^26^. To put the comparisons on a quantitative footing, we extracted the crossover frequency ω_c_ from the experimental data and scaled the computed crossover frequency to match the measured value. This adjustment was necessary for direct comparisons since the simulations are of coarse-grained representations and do not include explicit representations of the solvent molecules or solvent-mediated interactions. No other fitting or adjustments were made beyond this single parameter rescaling.

Comparisons of the computed and measured storage and loss moduli show that the computations based on the single-chain model systematically overestimate the moduli across the frequency range that was probed experimentally. While there is reasonable agreement with the loss moduli in the intermediate frequency range, the storage modulus computed using the single-chain model is a systematic overestimate vis-à-vis the measurements. These deviations were surprising given the excellent agreement between measured and computed phase diagrams for A1-LCD and numerous designed variants thereof ^47^. This prompted us to appreciate that the measurements, which construct complex shear moduli from autocorrelation functions of the motions of beads within condensates, investigate the material properties of the dense phase as a whole and not just the contributions of individual chains. Furthermore, the concentrations of polymers within dense phases have been measured to be above the overlap concentration with the measured concentrations corresponding to the semidilute regime ^47^. In this regime, the differences between intra- and inter-chain interactions vanish, and chain segments, referred to as thermal blobs ^61^, move under the influence of inter-segment interactions, which may be within the same chain but are more likely to be between different chains. Additionally, dense phases may be viewed as percolated networks ^4,36,37,51^, and hence we reasoned that the network of chains making up the dense phase can be treated as a single chain. This leads to a collective model for Zimm matrices.

### B. The collective model

In the collective model, the nodes are individual chains within the dense phase (Fig. 4a). For a pair of molecules, an edge is established if at least one pair of residues are nearest neighbors on the cubic lattice, which we define as belonging to any of the 26 sites that surround a given lattice site. The graph constructed in this way is undirected, meaning that the edges are symmetric. In the dense phase, the number of inter-chain contacts per chain is at least an order of magnitude larger than the number of intra-chain contacts per chain (Fig. 4b). This is consistent with previous observations showing that individual chains are more expanded in dense phases when compared to coexisting dilute phases ^47,62-66^, with the expansion enabling a higher degree of networking within dense phases ^47^. Using this approach, it was previously shown that the internal organization of condensates formed by A1-LCD and other molecules corresponds to a small-world network ^47^. Given this observation, the coarse-graining procedure used to generate Zimm matrices for the collective model, whereby individual chains are replaced by a node (Fig. 4c), has precedent in the literature. Specifically, Song et al., used block renormalization to demonstrate that complex networks, including those with small-world topology, have self-similar, scale-free properties ^67^. This justifies our coarse-graining whereby each node is now a single chain as opposed to being a single bead. Unlike the sparsity of the Zimm matrix observed for the single-chain model (Fig. 3b), the Zimm matrix for the collective model is considerably more dense on average. The hub-and-spoke nature of the internal organization within dense phases gives rise to Zimm matrices that are differently dense or sparse (Fig. 4d) ^47^.

**Fig. 4:**
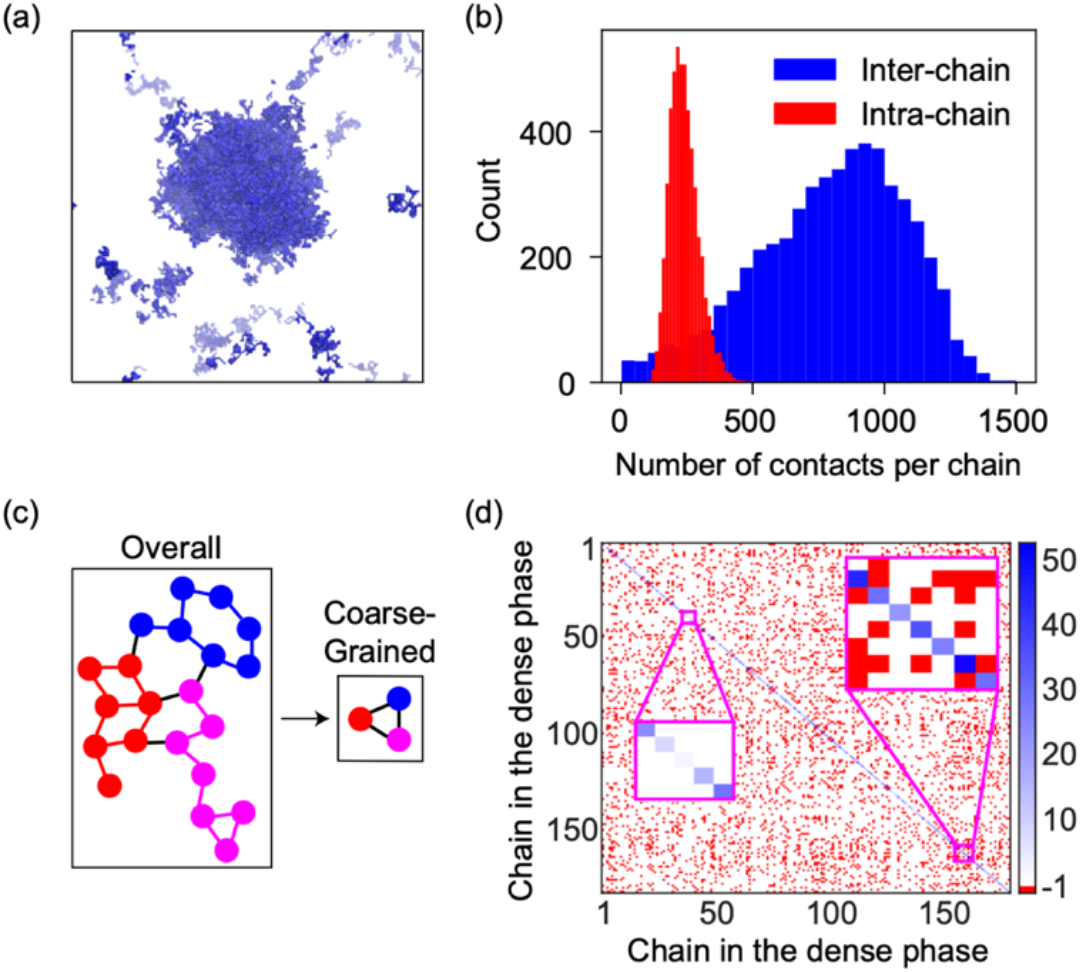
In the collective model, the entire polymer network is treated as a single chain. (a) We analyze dense phases from LaSSI simulations of WT^+NLS^. (b) Chains in the dense phase have a high likelihood of forming inter-chain interactions ^47^. (c) A coarse-grained model of the dense-phase network was developed by treating the entire network as a single chain. Each chain is treated as a node. An undirected edge between nodes indicates that at least one pair of residues between the chains forms a contact. (d) The Zimm matrix corresponding to the dense phase in panel (a). The insets indicate regions with different sparsity.

We compared the moduli, computed using the collective model to the measured moduli, using the same rescaling of frequencies that was used in the comparison of the single-chain model (Fig. 5a). Inclusion of the inter-chain contacts and treating the network of chains as a single chain leads to almost perfect agreement between the measured and computed loss moduli and good agreement across the intermediate- and high-frequency ranges for the storage moduli. This suggests that the actual network is more compliant than is anticipated by the single-chain model, meaning that the network can more readily deform in response to internal stresses ^19^. Intermolecular interactions in the semidilute regime help lubricate the motions of molecules and segments. This point is made by comparing the relaxation modulus computed using the single-chain versus collective models (Fig. 5b). We note that in the collective model there is a shortening of the relaxation times when compared to the distribution computed using the single-chain model. The network is more compliant when compared to what we infer by monitoring the relaxation modes of a single chain under the influence of intra-chain interactions alone.

**Fig. 5:**
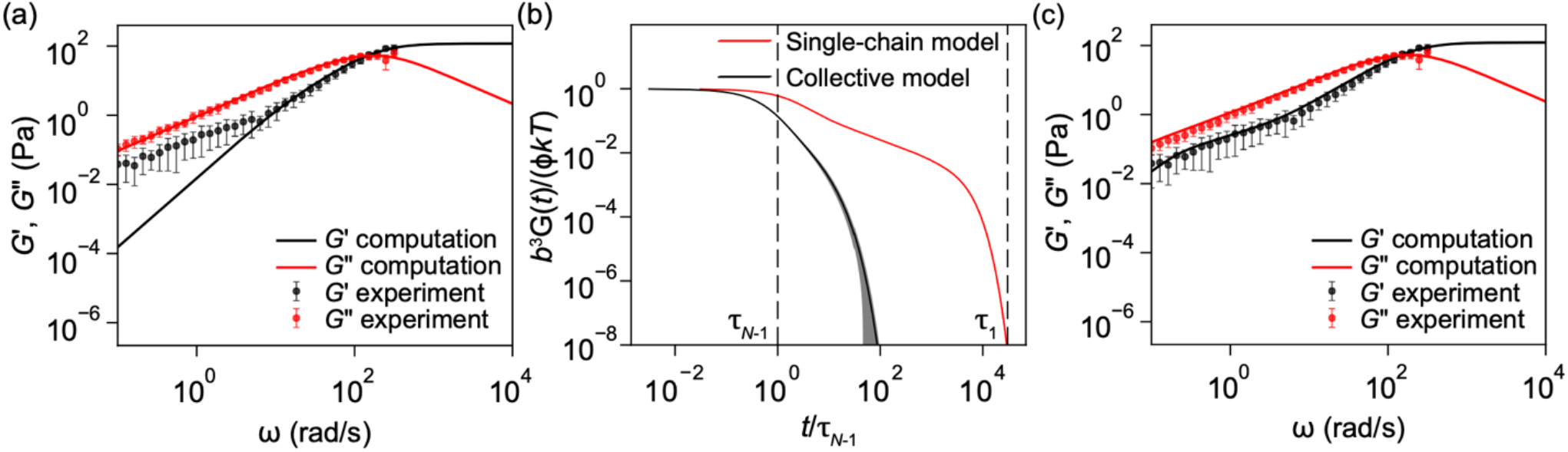
Performance of the collective model, the computed relaxation modulus, and a mixture model. (a) The storage and loss moduli computed using the collective model are plotted alongside the experimental values for WT^+NLS^. (b) The relaxation modulus shown here is computed using both the single-chain and collective models. The collective model leads to a shortening of the relaxation times as the network is less stiff and more compliant. In (a) and (b), 30 snapshots were analyzed over 3 replicates. The error bands indicate the standard deviation. (c) A linear combination of the moduli computed using the single-chain and collective models reproduces the experimental values with a coefficient of ≈ 0.88 for the collective model.

### C. The mixture model

Returning to the comparison between computed and measured moduli, we note that the storage moduli in the measurements deviate from the computed ones at low frequencies. This was initially viewed as being likely due to noise in the microrheology measurements ^26^. However, closer scrutiny rules this out as a valid explanation. Instead, it appears that while the collective model does a good job of recapitulating the elasticity of the network at intermediate and high frequencies, it underestimates the elasticity at low frequencies. Individual polymers are not coarse-grained beads freely moving around one another. While this coarse-graining is valid for describing the network, it appears that on longer timescales the motions combine the attributes of the collective and single-chain model. This proposal is based on the observation that the measured values for storage moduli lie between those of the single-chain and collective models at low frequencies. Accordingly, we asked if a mixture model can be parameterized to capture the totality of the measured frequency dependence of storage and loss moduli. In the mixture model, the measured storage modulus is postulated to be a linear combination of the storage moduli computed from the collective (*cm*) and single-chain (*sm*) models:

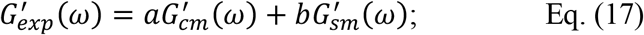

This is subject to the constraint that the coefficients *a* and *b* are positive scalars, and that *a* + *b* = 1. To determine the values of the coefficients, we performed an optimization using gradient descent. The learning rate, *γ*, was 0.06, and we iterated up to 2000 times or until convergence, with a numerical tolerance of 10^−6^. All data were first normalized to ensure the reliability of the procedure. This was done by scaling and centering the data such that they have a mean of zero and a standard deviation of one. For each iteration, the error was computed as the difference between the linear combination as defined above and the normalized target (experimental) data. The coefficient *a* was updated according to:

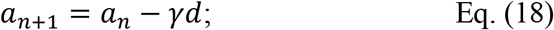

Here, *d* is the arithmetic mean of the error multiplied by the normalized test (*cm*) data. The coefficient *a* was then reset to the original scale by multiplying the value by the ratio of the standard deviation of the unnormalized experimental data to the standard deviation of the data from the collective model. Using this procedure, we obtain *a* ≈ 0.88, and *b* = 1 − *a*.

The optimized values of the coefficients *a* and *b* were used in Eq. (17) to compute the storage and loss moduli, and the frequency dependence was compared to the measured moduli. Note that the estimation of the coefficients did not use the measured loss moduli as a target. The mixture model generates profiles that are in very good agreement with the measurements for both the storage and loss moduli (Fig. 5c). The physical implication is that viscoelastic moduli of dense phases are influenced by a combination of the inter-chain contacts that give rise to a percolated, small-world network and the rigidity that arises from the segments being part of a polymer. Accordingly, threadlike molecules, to use the phrasing of Rouse ^25^, form networks through a collection of inter-chain contacts that make and break across a range of timescales. The long-time motions are hindered by the internal friction of a single chain ^62,68^, and this diminishes the compliance of the network. As a result, both inter-chain interactions, which are dominant in semidilute solutions, and intra-chain contacts, which are captured in the single-chain model, jointly, albeit disproportionately, contribute to the totality of the frequency-dependent viscoelasticity of A1-LCD condensates. The precise parsing of the contributions from the single-chain versus collective models are likely to be system-specific. For the systems studied to this point, the parsing appears to be equivalent to that of the WT^+NLS^ system, although this will need extensive scrutiny across a wide range of systems.

Before concluding this section, we note the following: Below the crossover frequency, the storage modulus is smaller than the loss modulus. However, this does not imply that the dense phases are purely viscous materials that, via physical aging such as a glass transition ^69^ or conversion to solids ^70^, become dominantly elastic ^58^. Such binary characterizations of early-time condensates as being viscous fluids that age and convert to elastic solids glosses over the fact that even on timescales when the loss modulus is larger than the storage modulus, the two moduli are within an order of magnitude of one another (Fig. 5c) and the materials are viscoelastic. This negates the characterization of liquid-like condensates being purely viscous Newtonian fluids ^16,32,69,71,72^. Instead, the finite storage modulus gives dense phases a network-like structure ^39,47,58^ and this will have a direct influence on the mechanical forces transduced, sensed, and exerted by condensates ^24,73-78^.

## V. CONTINUOUS DISTRIBUTION OF RELAXATION TIMES

Measured moduli of condensates suggest that these materials belong to the same class as Maxwell fluids ^10,12,26,69^. A Maxwell element features a spring and a dashpot in series ^57,79^. In coarse-grained descriptions based on appropriate constitutive equations, a Maxwell fluid with a single element will generate a frequency-dependent profile akin to what has been measured for A1-LCD-based systems ^57,69,80^. This has been taken to mean that condensates can be reduced to a single Maxwell element and that the presence of a single crossover frequency implies that condensates are defined by a single relaxation time ^69^. However, as shown by Rouse ^25^, the number of relaxation modes is governed by the number of non-zero eigenvalues of the Zimm matrix. Further, each mode can be thought of as a Maxwell element ^25^. Accordingly, if dilute or semidilute polymer solutions are dominantly viscous at long times, then they are best described as generalized Maxwell fluids ^58^, whereby Maxwell elements, one per each mode, are assembled in parallel ^81^. While the continuous relaxation spectrum, *h*(τ), cannot be measured directly, it can be extracted by solving an inverse problem because it is related to the storage and loss moduli via:

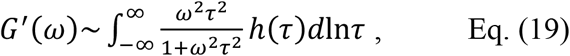

and

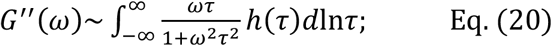

Note that the continuous relaxation spectrum is related to the discrete spectrum by

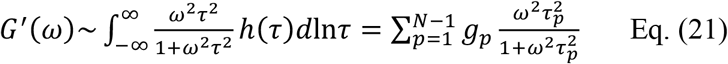

and

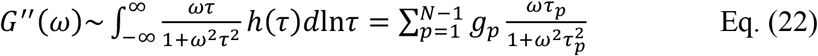

For *N* monomers or chains there are (*N* − 1) Maxwell modes giving finite relaxation times, and {*g*_*p*_, τ_*p*_} describe the weights and relaxation times, respectively, for the *p*-th mode. We do not have *a priori* knowledge of the weights, but we have the measured moduli, and we also have the computed moduli that match the measurements via the mixture model. Accordingly, we can use either as joint inputs and solve for *h*(τ). This a non-trivial inverse problem, and we solve it using a nonlinear Tikhonov regularization method ^82^. This method makes the substitution that *h*(τ) = exp(*H*(τ)S so that *h*(τ) > 0. We minimize the cost function given by:

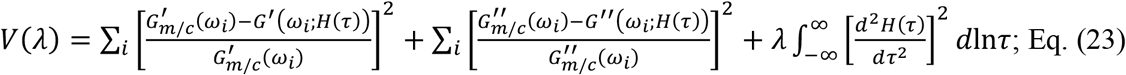

Here, *m*/*c* implies measured (*m*) or computed (*c*) moduli, and we use one or the other, but not both. In the current setup, we use the measured moduli. Eq. (23) can be rewritten compactly as *V*(λ) = σ^2^ + λ*η*^2^. The first term, σ^2^, denotes the mean-square error between the measured or computed moduli, 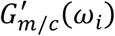 and 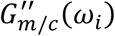 and the moduli inferred from the optimization procedure, *G*^′^(*ω*_*i*_; *H*(τ)S and *G*^″^ (*ω*_*i*_; *H*(τ)S, respectively, for the *i*-th frequency. Inferred moduli appear in the optimization because we are estimated the relaxation spectra whilst minimizing the deviations from the measured moduli. The second term consists of the norm of the curvature of *H*(τ) and the regularization parameter, λ, which controls the smoothness of *H*(τ). To obtain the continuous relaxation spectrum, we used the approach developed by Takeh and Shanbhag ^83^ that first uses a least-squares method to find the *H*(τ) while minimizing the cost function, and then determines the optimal λ from the so-called L-curve in the log-log plot of *η* vs. σ. We then compute *h*(τ) = exp(*H*(τ)S. As a consistency check, we used the computed *h*(τ) to solve the forward problem and extract the measured moduli from the estimated relaxation spectra.

Fig. 6 shows the relaxation spectra that are derived from the measured moduli. The relaxation time spectrum, *h*(τ), shows a broad, continuous distribution for the WT^+NLS^ and allY systems. Accordingly, we designate the condensates formed by these systems as being generalized Maxwell fluids featuring system-specific distributions of relaxation modes. Each relaxation spectrum has a distinct peak that corresponds to the system-specific crossover time in the dynamical moduli. Note that this crossover time shifts to longer values for the allY system compared to the WT^+NLS^, and this is consistent with the downshift of the crossover frequency. The upshot of our analysis is that measured moduli can be used to extract the spectrum of relaxation modes for condensates. This highlights the superiority of direct measurements of dynamical moduli over measurements of single-chain properties within condensates. This does not need the use of optical traps, although the methodology affords several advantages. Instead, light scattering or particle tracking measurements can be adapted to quantify complex shear moduli ^26,84,85^, and these measured moduli can be used to infer the relaxation spectra.

**Fig. 6:**
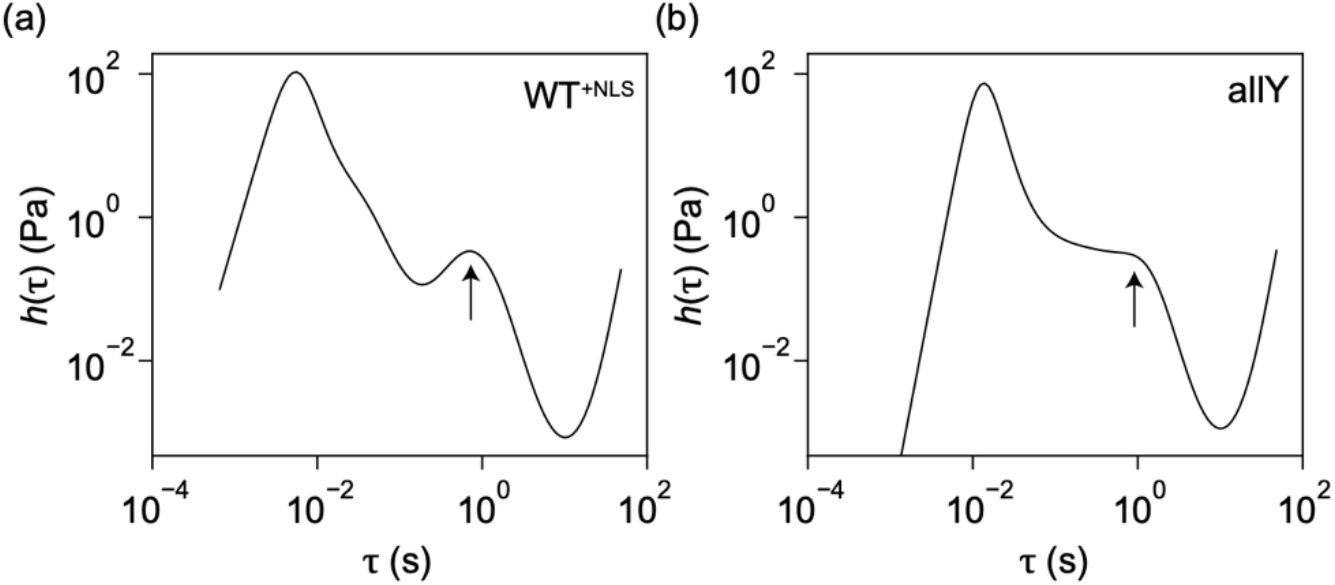
A continuous spectrum of relaxation times indicates that a generalized Maxwell model best describes dense phases of PLCDs. (a) We calculated the spectrum for the WT^+NLS^ from the experimentally measured moduli in Fig. 5a following a nonlinear regularization strategy. As a consistency check, we solve the forward problem exactly from the knowledge of *h*(τ), thereby recovering the storage and loss moduli. (b) We also compute the spectrum from the experimentally measured moduli for the allY variant following the same approach. The arrow in each panel indicates the peak in the relaxation time spectrum corresponding to the crossover in the dynamical moduli.

## VII. DISCUSSION

In this work, we have provided details of a recently introduced generalization of the Rouse model that incorporates different types of Zimm matrices whose eigenvalues are used to enable direct computations of viscoelastic moduli of dense phases formed by PLCDs. The Zimm matrices are computed using a graph-theoretic representation of lattice-based MMC simulations of condensate-forming systems. Two types of Zimm matrices were constructed: the single-chain model that accounts only for intra-chain contacts whereas the collective model accounts for inter-chain contacts. Overall, the collective model generates better agreement with the measured moduli. However, this model is not perfect, and a mixture of the two models with dominant contributions from the collective model explains the totality of measured, frequency-dependent moduli. The collective model, which is in accord with the dense phase being a semidilute solution ^47^, generates a compliant network. This model conceptualizes condensates as a collection of blob-sized segments that interact with one another to generate a percolated network. However, the motions of the segments are not unhindered, and the long-time dynamics require a proper accounting of the polymeric nature, as evidenced by the need for a mixture model to account for the totality of the frequency dependence of storage and loss moduli (Fig. 5c). At long timescales the collective model overestimates the compliance of the network, and the slithering of single chains and the sparsity of intra-chain contacts needs to be accounted for to generate the correct, frequency-dependent moduli as measured using microrheology.

Building on the foundational work of Rouse ^25^ and aided by the analysis of MMC simulations of two-phase systems, we demonstrate explicitly that the presence of a single crossover frequency is not sufficient to assert that there is a single relaxation time within a condensate ^69^. Instead, the computations show that there is a spectrum of relaxation times. These are continuous spectra. Accordingly, given the congruence between measured moduli and expectations for a Maxwell fluid, we propose that the dense phases may be viewed as generalized Maxwell fluid comprising a collection of Maxwell elements being assembled in parallel ^25,58^. This contrasts with there being a single Maxwell element that describes the entire system. Accordingly, condensates that are dominantly viscous are best described as generalized Maxwell fluids that are viscoelastic. The latter point is noteworthy because the storage moduli cannot be ignored even at long times, and both the storage and loss moduli contribute to the dynamics of condensates across the entire frequency range.

Importantly, we showed how the relaxation spectra can be extracted by solving an inverse problem with the measured or computed complex moduli as inputs. These relaxation spectra affirm the continuous nature of relaxation modes that span several orders of magnitude, corroborating recent suggestions of there being “extreme dynamics” ^62^ within fluid-like condensates.

The methods introduced here show how dynamical moduli of dominantly viscous, albeit truly viscoelastic materials can be computed from simulations of two-phase systems formed by disordered proteins, which are becoming increasingly tractable both at the coarse-grained and fine-grained levels ^28,40,47,62,64,66,72,86-91^. However, for MMC simulations, we do not have information about timescales, nor do we have access to hydrodynamic interactions that are mediated by solvent contributions. While molecular dynamics simulations do not have such issues, the timescales that are accessible cannot span the range of relaxation modes that define dynamics within condensates ^62,72,91^. Accordingly, analysis of simulations, without any input from experiments, can be useful for extracting robust insights that enable relative comparisons, but the computed values of absolute moduli will be unreliable. However, knowledge of the crossover frequency is more than sufficient to put the computed moduli on realistic time and energy scales. Importantly, the computations in concert with microrheology measurements, deployed across a range of systems, have the promise of enhancing our understanding of the connections between driving forces for condensate formation and the material properties of condensates.

Finally, there is considerable interest in the physical aging of condensates ^22,26,27,58,69,70,92-98^. If cages form within condensates ^99,100^ or there is a time-dependent increase in intra-condensate rigidity, then the lifetimes of stickers will likely increase ^58^, and this can contribute to physical aging. Such a process, whereby both the elastic and loss moduli can increase, and the crossover frequency becomes lower, would correspond to a glass transition that generates a network glass ^99,101^. Indeed, computations have shown that long-lived crosslinks, which can be irreversible on the timescales of simulations can arrest condensate coarsening ^97^ and generate core-shell structures even for single-component systems ^95^. The latter finding takes on special significance given recent studies showing that the generation of compositionally distinct condensates *in vitro* and in cells can be under dynamical control, whereby differences in sticker lifetimes can engender compositionally distinct protein-RNA condensates that coexist with one another ^102,103^. Dynamical arrest due to competing protein-DNA and protein-RNA interactions has also found to be operative in generating dissipative structures in active condensates ^104^.

An alternative to rigidity transitions or glass transitions mediated by enhanced sticker lifetimes ^58^ is the conversion to the globally stable ground state, which is an equilibrium solid defined by a disorder-to-order transitions ^26,94,98^. This is not the same as aging due to enhancements of the lifetimes of inter-sticker interactions ^58^. Instead, for PLCDs, it was shown that weakening sticker crosslinks and replacing naturally occurring spacers with those that enhance the preference for chain solvation substantially weakens the driving forces for forming disordered fluid phases ^26,29,47^. However, these mutations lower the barrier for conversion to non-fibrillar solids ^26^. These solids feature increased beta-sheet contents and have semi-crystalline morphologies ^26^. Active microrheology, specifically creep tests, show that the equilibrium solids have the elastic responses of Kelvin-Voigt solids ^26^. An interesting challenge for MMC simulations is to model the changes in material properties that result via conversions of metastable fluids to globally stable solids.

## ACKNOWLEDGMENTS

This work was supported by the St. Jude Research collaborative on the Biology and Biophysics of RNP granules (P.R.B and R.V.P), the US Air Force Office of Scientific Research (grant FA9550-20-1-0241 to R.V.P), and the US National Institutes of Health (R01NS121114 to R.V.P). We thank Mina Farag for the simulations of the A1-LCD system and our collaborators, Ibraheem Alshareedah and Tanja Mittag, for helpful discussions. The code for computing Zimm matrices is available on GitHub at https://github.com/Pappulab/material_properties.

